# NMI induces chemokine release and recruits neutrophils through the activation of NF-κB pathway

**DOI:** 10.1101/2024.09.13.612823

**Authors:** Zhenxing Chen, Yongjie Yao, Yuzhou Peng, Zhuangfeng Weng, Yingfang Liu, Na Xu

## Abstract

Neutrophils are essential components of the innate immune system, playing a pivotal role in immune responses. These cells rapidly migrate to sites of inflammation or tissue injury, facilitating pathogen clearance and tissue repair. The chemotactic signaling network regulating neutrophil recruitment is complex and not fully understood, particularly regarding damage-associated molecular patterns (DAMPs). In our previous research, we identified NMI as a DAMP that activates dendritic cells and macrophages, amplifying inflammatory responses and contributing to both acute and chronic inflammation. In this study, we investigated the role of NMI in neutrophil recruitment. We purified and characterized a recombinant murine NMI protein, ensuring endotoxin removal while preserving biological activity. In vivo experiments demonstrated that NMI enhances neutrophil recruitment in both a murine air pouch model and an acute peritonitis model, mediated by macrophage-derived chemokines. In vitro assays revealed a concentration-dependent increase in neutrophil migration induced by NMI, facilitated by chemokine secretion and subsequent migration through the CXCR2 receptor. Importantly, we established that NMI activates chemokine expression via the NF-κB signaling pathway. These findings provide insights into the mechanisms of NMI-induced neutrophil migration, enhancing our understanding of neutrophil recruitment during inflammation.

## Introduction

Neutrophils, the most abundant type of white blood cells in human circulation, play a crucial role in the body’s immune response. Their recruitment and function are essential for directing inflammatory cells to sites of infection, where they can combat invading pathogens while minimizing damage to host tissues or triggering autoimmune reactions [1]. However, excessive neutrophil infiltration can have detrimental effects, exacerbating inflammation and tissue damage, and potentially contributing to the development of various diseases, such as acute pancreatitis [2], alcoholic liver injury [3], and autoimmune disorders [4-6]. In recent years, there has been growing interest in understanding the underlying mechanisms that regulate neutrophil recruitment.

Neutrophil recruitment is the process by which neutrophils migrate from the circulatory system to the site of inflammation, directed by various chemotactic signals. Typically, chemokines serve as the primary chemotactic signals [7]. Neutrophils, macrophages, mast cells, endothelial cells, etc. can secrete a large number of chemokines in response to proinflammatory cytokines (e.g., IL-1β, TNF, IFNγ, IL-17). Besides, pathogen-associated molecular patterns (PAMPs), and damage-associated molecular patterns (DAMPs) are also inducers of chemokine administration [8]. In fact, The secretion of chemokines is regulated through a complex process, with the NF-κB signaling pathway playing a crucial role in modulating chemokine release. [9].

DAMPs are particularly important as chemoinducers of neutrophil recruitment. After being released into the extracellular space, DAMPs activate endothelial cells, macrophages, and mast cells, leading to the release of a large number of proinflammatory cytokines and chemokines [10]. Furthermore, DAMPs can stimulate the NF-κB signaling pathway via Toll-Like Receptors (TLRs) or Receptor for Advanced Glycation End-products (RAGE), including molecules such as High Mobility Group Box 1 (HMGB1), S100A8/9, and interferon-inducible protein 35 family proteins (IFP35) [11-13]. However, the precise role of DAMPs in neutrophil recruitment remains to be fully elucidated.

N-myc-interactor (NMI) and IFP35 are two members of the IFP35 family of proteins. In a previous study, we found that NMI and IFP35 act as DAMPs released after cell death to activate the NF-κB signaling pathway extracellularly via TLR4, which activates the release of pro-inflammatory cytokines from macrophages and exacerbates inflammatory responses. Furthermore, our research has shown that knockout of *nmi* and *ifp35* in mice increased their survival in an LPS-induced septic shock model, and that NMI and IFP35 are released in large quantities during cellular injury and pathogen invasion [14]. These released molecules can also serve as potential biomarkers or therapeutic targets for a variety of inflammatory diseases, such as multiple sclerosis and lung injury [15, 16]. Their involvement in these pathological processes underscores the importance of further investigating NMI and IFP35 in the context of inflammatory disease mechanisms and therapeutic interventions.

In our previous study on APAP-induced liver injury model, we observed a significant reduction in neutrophil infiltration in the liver of ifp35 knockout mice after liver injury [17]. This finding led us to hypothesize that NMI might influence the recruitment of neutrophils during inflammatory responses triggered by infection or injury. In the present study, we found that NMI can recruit large numbers of neutrophils in mice, this process proceeds in part through macrophages. In addition, NMI can lead to the expression and release of chemokines from neutrophils by activating the NF-κB signaling pathway and induce neutrophil migration through the CXCR2 receptor. Our findings provide new insights into the mechanisms underlying neutrophils migration during inflammatory responses.

## Materials and methods

### Animals

8-10-week-old male wild-type C57BL/6 mice were purchased from SPF (Beijing) biotechnology Co. Ltd. All mice were maintained in a specific pathogen-free facility at the Experimental Animal Center of Sun Yat-Sen University. All animal experiments were performed according to the Ministry of Health national guidelines for housing and care of laboratory animals and approved by the Institutional Animal Care and Use Committee of Sun Yat-Sen University (SYSU-IACUC.MED-2022-B083), and all experiments conform to the relevant regulatory standards.

### Mouse air pouch model

The mouse air pouch model was created according to the previous studies with minor modifications [18]. To generate the air pouch, mice were anesthetized with isoflurane, and 6ml of filtered sterile air was subcutaneously injected in the back on Day 0, followed by an injection of 3ml of sterile air on Day 3 for re-expansion of the air pouch. To evaluate neutrophil recruitment, on day 6, mice were killed 6 h after injection of sterile PBS or LPS (1 μg/mL) or NMI (100 μg/mL) into the air pouch, the LPS was used as the positive control. Thereafter, the air pouch lavage was collected and the number of emigrated neutrophils was quantitated by FACS analysis by staining for Ly6G, CD45, CD11b.

### Acute peritonitis model

Thioglycollate-induced peritonitis in *wild-type, nmi*^-/-^ mice was performed as described previously [19]. Alternatively, wild-type mice were injected i.p. with 100 μg of NMI recombinant protein. For macrophage deletion studies, 1 day before the injection of 100 μg NMI recombinant protein, 100 μg of clodronate liposomes were administered i.v. to mice. Control mice were treated with the same volume of PBS. To evaluate peritoneal neutrophil recruitment, mice were killed at 4 hours following injection of thioglycollate or NMI recombinant protein. Thereafter, the peritoneal lavage was collected and the number of emigrated neutrophils was quantitated by FACS analysis by staining for Ly6G, CD45, CD11b.

### Induced differentiation and stimulation of bone marrows-derived macrophages cells (BMDMs)

*Wild-type* mice were sacrificed and bone marrow cells from the leg bones and tibia were isolated. Bone marrow cells were cultured in DMEM medium containing 1% penicillin and streptomycin and 10% FBS. The cell density was adjusted to 1×10^6^ cells /mL and added to 6-well plates. Added M-CSF (20 ng / mL) and then incubated at 37°C in a 5% CO_2_ incubator. Every 2 days, changed 1/2 culture medium, replenish 1/2 M-CSF. On 7th day, and the obtained primary macrophages were incubated with or without LPS (100 ng/mL), NMI (10 µg/ml), for 4 hours. Macrophages were collected for subsequent qRT-PCR assay.

### Isolation and stimulation of Bone Marrow Neutrophils (BMNs)

Mouse bone marrow was collected from femur and tibia bones. BMNs were isolated using Histopaque-1119 (1191-100ML, Sigma) and Histopaque-1077 (1077-100ML, Sigma) according to the manufacturer’s instructions. The purity of BMNs populations was greater than 93% as determined by flow cytometry after staining with anti-Ly6G-APC (B357443,BioLegend), and the viability was greater than 99% as determined by trypan blue staining. For transwell migration assay, the primary neutrophils were pretreated with the concentration of 1 μM CFSE (21888, Sigma), the cells can be observed to light green (Green Fluorescent). Besides, the primary neutrophils were incubated with or without LPS (100 ng/mL), NMI (10 µg/ml), for 2 hours, neutrophils and supernatants were collected for subsequent qRT-PCR, western blotting and ELISA assays. In the inhibition group, primary neutrophils were pretreated with inhibitors for 30 min. The inhibitors and their inhibition sites: TAK242 (S7455, Selleck): inhibitor of TLR4, Repertaxin (S8640, Selleck): inhibitor of CXCR2, AMD3100 (S8030, Selleck): inhibitor of CXCR4, QNZ (S4902, Selleck): inhibitor of NF-κB signaling pathway.

### Transwell migration assay

Chemotaxis of isolated neutrophils was assayed with a modified Boyden chamber with 8.0 µm pores (Corning, USA). Mouse bone marrow neutrophils were rested in RPMI 1640 medium for 1 hour. After resting, the neutrophils were plated into the upper compartment of the transwell, and RPMI 1640 medium, with or without chemoattractants, was added to the lower wells beneath the polycarbonate membrane, fMLP added at the concentration of 100 ng/mL as positive control, and added NMI recombinant protein at a concentration of 10 µg/mL as a stimulation group. After 2 hours of stimulation, the number of cells in the lower chamber was observed under a fluorescence microscope. Resulting statistics: number of differentially migrating cells relative to the PBS treated group (number of cells in the lower chamber in the induced group/number of cells in the lower chamber in the uninduced group).

### Under-Agarose cell migration assay

The under-agarose cell migration assay was performed as previously reported [20]. After incubation at 37°, with 5% (v/v) CO_2_ for 2 hours, the chemotaxis distance was used to determine the chemotaxis of neutrophils. The images were captured by an Eclipse Ti-E Inverted Microscope (Nikon) and NIS-Elements AR analysis software was used for data collection.

### Quantitative Real-Time Polymerase Chain Reaction (qRT-PCR) Assays

The total RNA was isolated from neutrophils using TRIzol. cDNA was synthesized using the ReverTra Ace qPCR RT Kit (R423-01; Vazyme). Quantitative PCR was performed using SYBR Green PCR Master Mix (ZA352-101; Gen Star) with a QuantStudio™ 5 System, and the relative quantitative analysis of the gene was performed using the 2^-ΔΔ^CT method. The primers utilized for amplification of the cDNA were as follows (forward, reverse): GAPDH, 5’-AGCCT CGTCCCGTAGACAA-3’ and 5’-AATCTCCACTTTGCCACTGC-3’; CXCL1, 5’ - CCAAACCGAAGTCATAGCCAC-3’ and 5’-TCCGTTACTTGGGGACACCT-3’; CXCL2, 5’-TCCAAAAGATACTGAACAAAGGCAA-3’ and 5’-GCGAGGCACA TCAGGTACGA-3’; CCL3, 5’-CCAAGTCTTCTCAGCGCCA-3’ and 5’-CGGTT TCTCTTAGTCAGGAAAATGA-3’.

### Enzyme-Linked Immunosorbent Assays (ELISA)

The quantification of CXCL1 (abs520017-96T, Absin), CXCL2 (abs520018-96T, Absin) was performed using quantitative ELISA. Optical densities were measured at 450 nm using an ELISA-dedicated instrument. CXCL1 and CXCL2 concentrations were calculated using a standard curve. All measurements were performed in duplicate, and arithmetic averages were calculated.

### Expression and purification of NMI recombinant protein

Recombinant mouse His-NMI protein were expressed in *E*.*coli*. The plasmid of NMI was transformed into E. coli strain BL21(DE3). Cells were cultured in Luria-Bertani medium at 37 °C with 50μg/mL kanamycin. When the OD600 reached 0.8–1.0, the culture was induced by addition of isopropyl-β-D-thioglactosidase (11411446001, Sigma) to a final concentration of 0.3 mM for 20 hours at 16 °C. Cells were collected by centrifugation at 2500 g for 15 min. Pellets were resuspended in lysis buffer (25 mM Tris at pH 8.0, 400 mM NaCl, 30 mM imidazole) and lysed by sonication and the lysate was separated by centrifugation at 20000 g for 30 min, and the recovered supernatant was applied to a Ni-NTA affinity column(SA004100S, mart-lifesciences), followed by intensive washing with washing buffer (25 mM Tris at pH 8.0, 150 mM NaCl, 30 mM imidazole). Recombinant protein was eluted from the Ni-NTA affinity column using elution buffer (25 mM Tris pH 8.0, 100 mM NaCl, 500 mM imidazole) and further purified by Anion Exchange Chromatography Columns using buffer (A:25 mM Tris pH 8.0, 100 mM NaCl; B: 25 mM Tris pH 8.0, 1000 mM NaCl). For The final part of purification by gel filtration with a Superdex 200 column (GE Healthcare) using PBS buffer. The purity and integrity of recombinant proteins were verified by Coomassie blue staining after SDS–PAGE. Finally, endotoxin is removed using Pierce High-Capacity Endotoxin Removal Resin (88276, Thermo). Limulus amebocyte lysate assay was used to detect the potential endotoxin contamination.

### Determination of NMI Protein Activity

The experimental methodology employed follows the guidelines for utilizing the RAW-Blue™ cell line (Invivogen, USA). briefly, RAW Blue cells were categorized into several treatment groups: PBS (negative control), 100 ng/mL LPS (positive control), 10 μg/mL NMI, 10 μg/mL NMI combined with PMB, 10 μg/mL NMI combined with Trypsin, and 10 μg/mL NMI combined with both PMB and Trypsin. The resulting protein activity was quantified by measuring the SEAP levels at wavelengths between 620 and 655 nm. PMB was incorporated into the experimental design to mitigate the risk of endotoxin contamination in the recombinant proteins.

### Western Blot Analysis

The total protein of BMNs were extracted with the RIPA Lysis Buffer (P0013, Beyotime) containing protease and phosphatase inhibitors (P1045, Beyotime). Protein concentration was determined using the BCA assay. Equal concentrations of protein were separated on 10% SDS-PAGE gels and transferred onto NC membranes. Following blocking for 1 h at room temperature with 5% BSA, the membrane was incuated overnight with the primary antibody: anti-GAPDH (5174S, CST; Anti-mouse NF-kappaB p65 (T50034, Abmart); Anti-mouse Phospho-NF-kappaB p65 (Ser536) (93H1) (3033T, CST). After three washes, the membranes were incubated with fluorescence-conjugated anti-rabbit (7074S, Cell signaling) for 1 h at room temperature. Proteins were visualized using an ECL kit (Beyotime Biotechnology) and quantified using a Tanon 5500 imaging system (Tanon, China).

### RNA-seq and differential expressed gene dentification, annotation and enrichment analysis

Bone marrow derived neutrophils after stimulated in vitro were used for total RNA isolation with TRIzol (Invitrogen). RNA-Sequencing services were provided by BGI Genomics Co., Ltd. Shenzhen, China. Using R package DESeq2, differential expression analysis was performed on the count data and significant pathway by GO enrichment analysis.

### Statistical analysis

All data were analyzed using GraphPad Prism software (GraphPad10.1.2 Software). Data were analyzed using by one-way ANOVA. P < 0.5 was considered statistically significant for all experiments. All values are presented as themean ± SEM.

## Results

### 1. NMI recruits large numbers of neutrophils in mouse air pouch model

We previously observed that the knockout of *ifp35* led to a significant reduction in neutrophil infiltration within the APAP-induced liver injury model. To further explore whether NMI could induces neutrophil recruitment, we purified and characterized a recombinant murine NMI protein (Fig. S1) and established a mouse air pouch model based on established methodologies (Fig. 1a). Upon injecting the NMI recombinant protein into the air pouch of *wild-type* mice, we noted a marked Infiltration of inflammatory cells within a 6 hours timeframe (Fig. 1b). Notably, among these immune cells, a substantial proportion of neutrophils (CD11b^+^Ly6G^+^/CD45^+^%) were recruited to the air pouch in response to the NMI protein (Fig. 1c). These findings indicate that NMI stimulation can provoke robust immune responses and facilitate neutrophil infiltration in vivo.

**Fig. 1.**
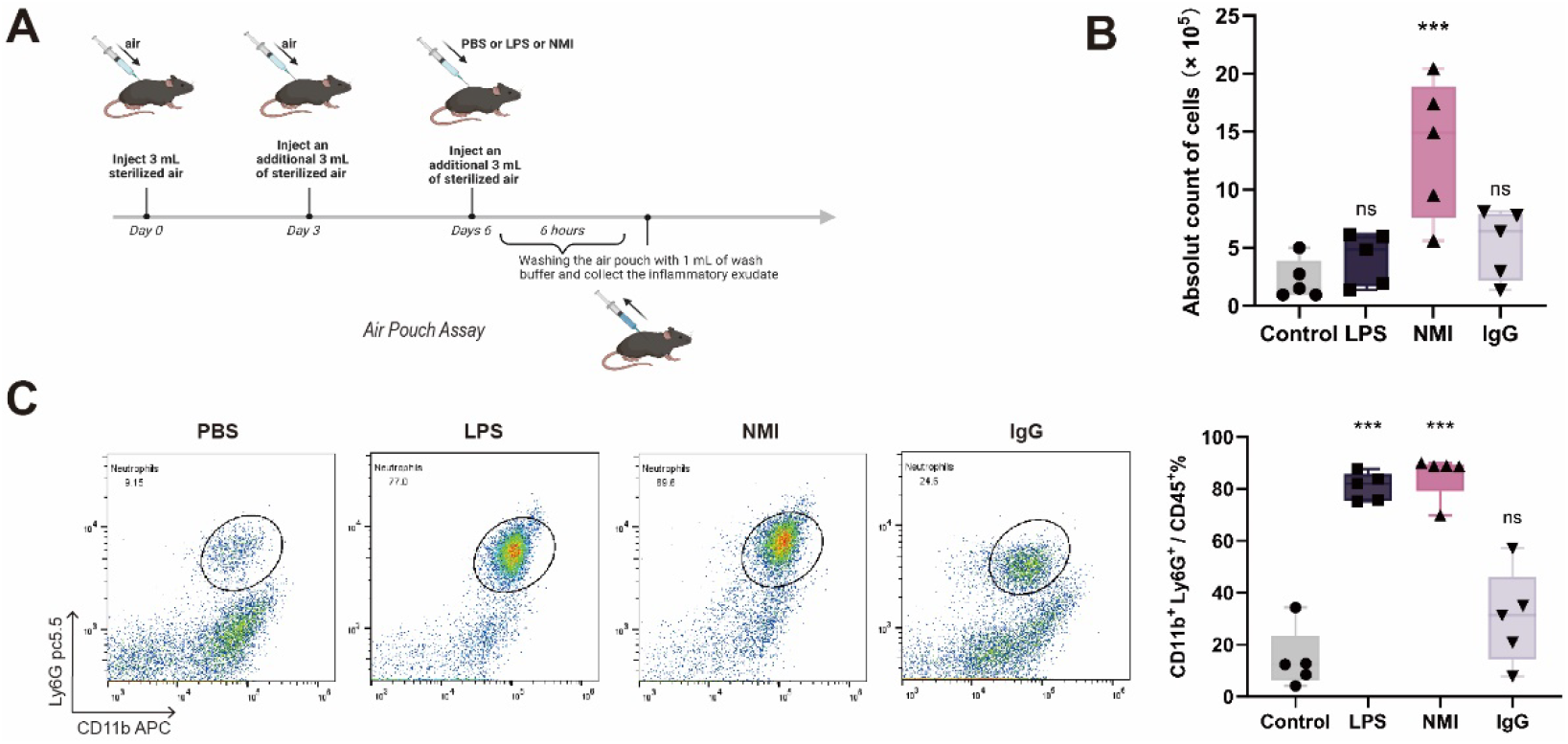
NMI induces neutrophil recruitment in mouse air pouch model. **a** Mouse air pouch model. **b** Flow analysis graphs and statistics of neutrophils in the air cavity lavage fluid. **c** Absolute counts of total cell density in airspace lavage fluid of mice. Statistical significance was determined using one-way ANOVA. (n=5, *P < 0.05, ** P < 0.01, *** P < 0.001)

### 2. In the mouse model of acute peritonitis, NMI recruits neutrophils through macrophages

To further investigate the mechanism by which NMI facilitates neutrophil recruitment, we developed a mouse model of acute peritonitis, as illustrated in Fig. 2a. Our observations revealed a substantial influx of neutrophils into the peritoneal cavity four hours post-intraperitoneal injection of recombinant NMI protein. Notably, knockout of the *nmi* gene did not impede neutrophil recruitment triggered by thioacetate injection in mice (Fig. 2b), suggesting that NMI may exert a direct effect in vivo. Given that macrophages serve as primary sentinel cells responsible for relaying chemotactic signals that drive neutrophil recruitment [21], we proceeded to stimulate bone marrow-derived macrophages (BMDMs) with NMI recombinant protein in vitro. As depicted in Fig. 2c-e, we observed a significant upregulation in the transcript levels of neutrophil recruitment-associated chemokines, including CXCL1, CXCL2, and CCL3, following NMI stimulation.To further elucidate the role of macrophages, we employed clodronate liposomes to selectively deplete macrophages in mice and subsequently assessed the neutrophil recruitment activity of NMI in the peritonitis model. Interestingly, the results indicated that neutrophil recruitment was only partially inhibited following macrophage depletion (Fig. 2f). These findings indicate that NMI significantly enhances neutrophil recruitment, partially through a mechanism that involves macrophages.

**Fig. 2.**
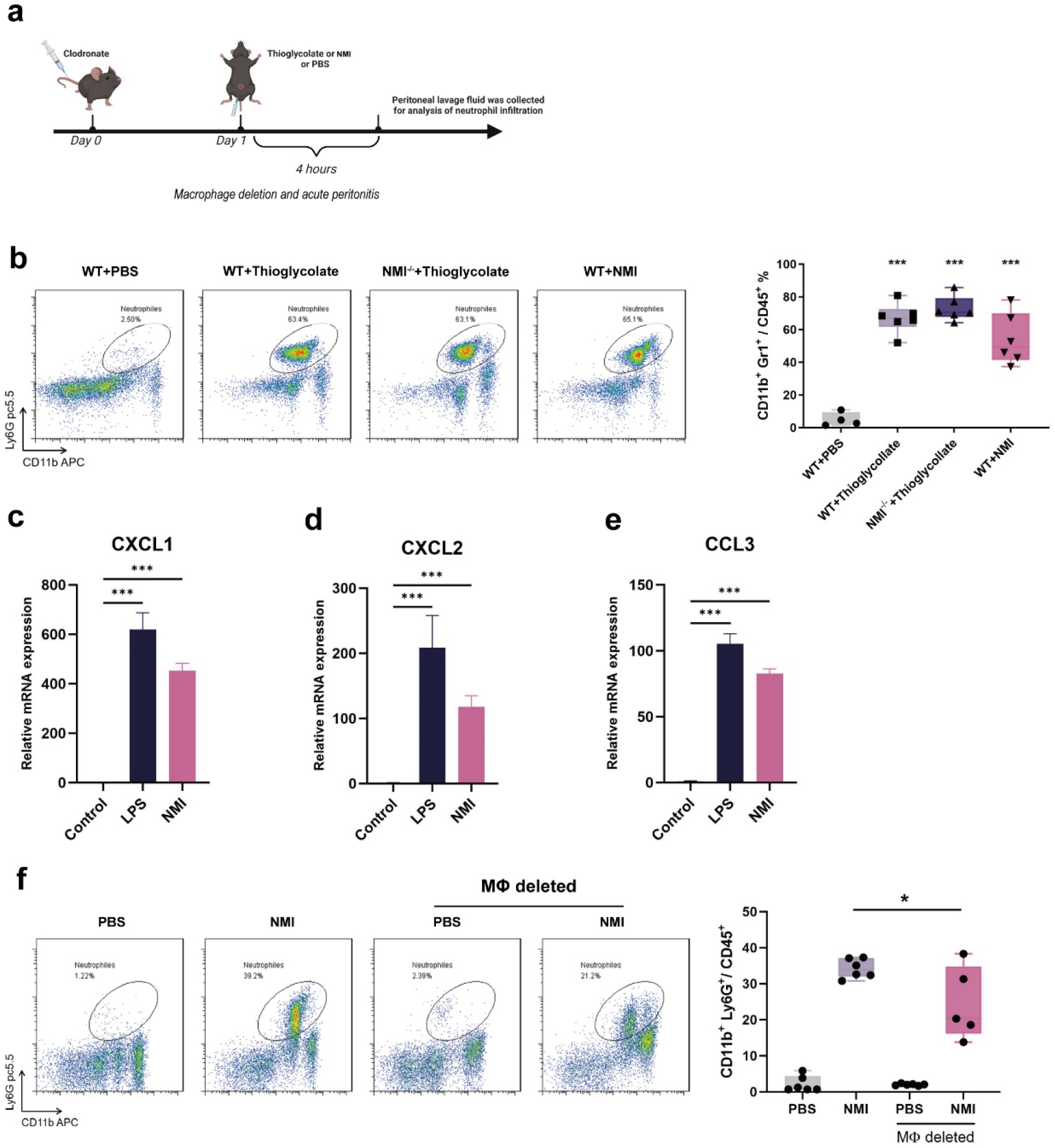
NMI recruits neutrophils in the acute peritonitis model. **a** Macrophage deletion and acute peritonitis model in mice. **b** Flow analysis plots and statistics of the peritoneal lavage fluid after 3 h of induction by injection of PBS, Thioglycollate, NMI (100 μg) into the peritoneal cavity of *wild type* mice and injection of Thioglycollate into the peritoneal cavity of *nmi*^-/-^ mice. **c-e** Differential expression of CXCL1, CXCL2, CCL3 in BMDM cells after 4 hours of NMI and PBS stimulation. (n=3). **f** FACS analysis of neutrophils recruited with PBS and 100 μg NMI in the presence of clodronate liposomes deleted from macrophages was plotted and counted (n = 5). All data are expressed as mean ± SEM. Statistical significance was determined using one-way ANOVA (*P<0.05, ** P<0.01, *** P<0.001).

### 3. NMI is chemically inducible to neutrophil chemotaxis

In order to further investigate whether NMI induces neutrophil migration in vitro, we also performed a transwell migration assay. We added different concentrations of NMI protein or 100 nM fMLP (a highly effective chemoattractant that induces neutrophil migration) to the lower wells of the polycarbonate membrane. After 2 hours of induction found that NMI induced neutrophil migration with a concentration dependent effect, and when the concentration of NMI was ≥5 μg/mL, neutrophil migration was significantly induced. Notably, NMI induced neutrophil migration at a concentration of 10 μg/mL comparable to 100 nM fMLP (Fig. 3a).

**Fig. 3.**
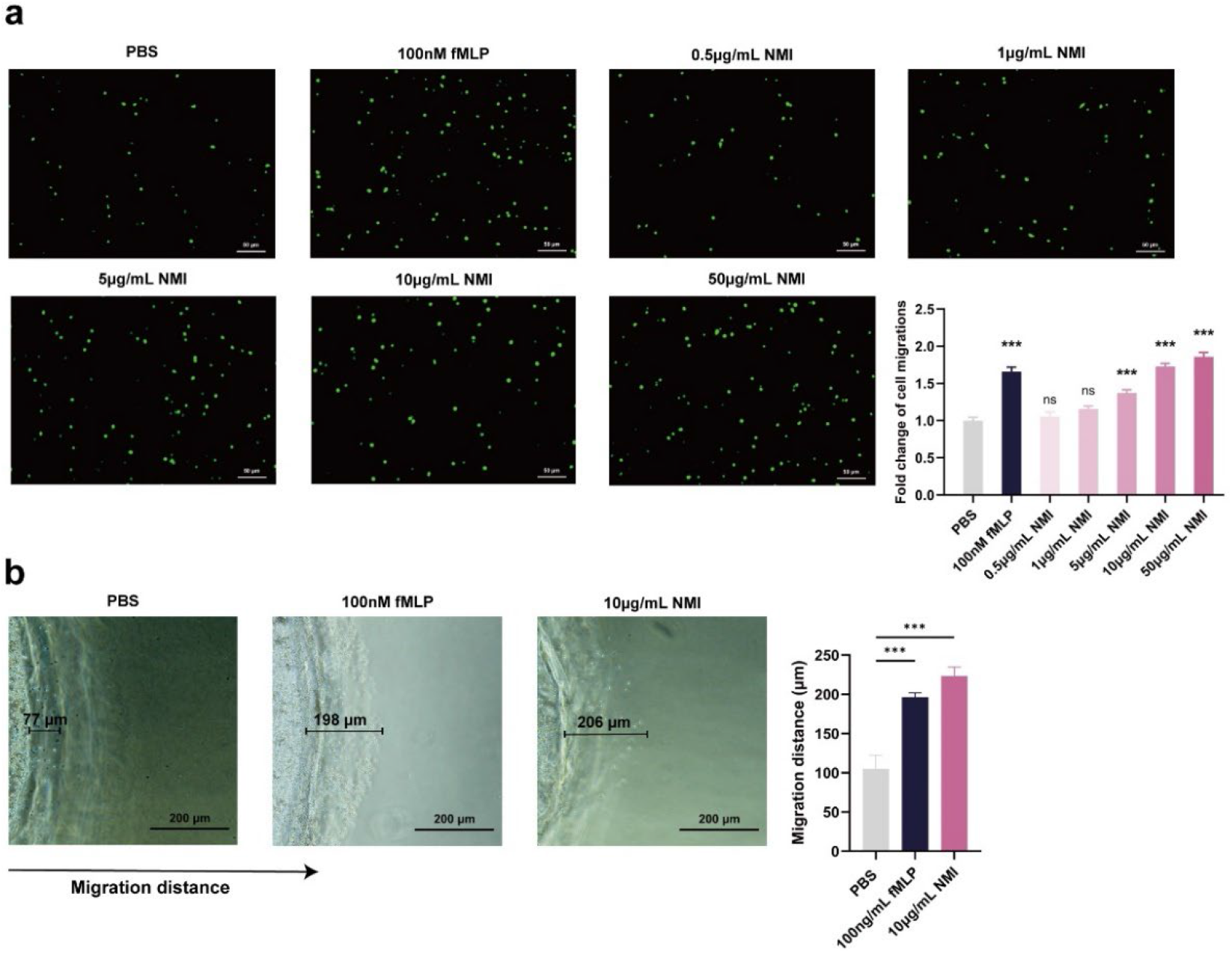
NMI is chemically inducible to neutrophil chemotaxis. **a** In vitro transwell migration assay. The fold change of neutrophils migrations during through transwell filters toward fMLP (100 nM), and different concentrations of NMI (0.5 μg/mL, 1 μg /mL, 5 μg/mL, 10 μg/mL, 50 μg/mL). **b** In vitro under-agarose cell-migration assay. The chemotaxis distance is represented by the distance of directional migrated neutrophils, neutrophils were treated with PBS 100 nM fMLP, 10 μg/mL NMI. All data are expressed as mean ± SEM. Statistical significance was determined using one-way ANOVA (n = 3, *P< 0.05, **P< 0.01, ***P< 0.001).

To further elucidate the impact of NMI on neutrophil chemotaxis, we employed the agarose-based cell migration assay. Notably, neutrophils migrated distances comparable to those induced by fMLP within a defined timeframe of 2 hours following the introduction of recombinant NMI protein (Fig. 3b). Collectively, these findings indicate that NMI directly enhances neutrophil migration in vitro, exhibiting a chemoinductive effect.

### 4. NMI induces neutrophil migration through the CXCR2 receptor

To explore the mechanisms underlying NMI-induced neutrophil migration in vitro, we first examined the direct interaction between NMI and its receptor TLR4. When we pretreated cells with TAK242, a TLR4 inhibitor, we observed that NMI protein only partially inhibited neutrophil migration (Fig. 4a). The visualization of neutrophil chemotaxis is presented in Fig. S2. Our earlier findings indicated that NMI stimulates macrophages to produce a substantial array of chemokines, which play a crucial role in driving neutrophil recruitment. It is noteworthy that neutrophils themselves are significant producers of chemokines, contributing to their own recruitment [22]. To further investigate this, we sought to inhibit the chemokine receptors CXCR2 and CXCR4, which play a crucial role in neutrophil migration [23]. We pretreated the cells with the CXCR2 inhibitor retaxin and the CXCR4 inhibitor AMD3100 for a duration of 30 minutes. Following this, we added PBS, fMLP (100 nM), and NMI (10 μg/mL) to induce neutrophil migration over a period of 2 hours. Surprisingly, the inhibition of CXCR2 resulted in a significant blockade of neutrophil migration, while the inhibition of CXCR4 showed only moderate effects (Fig. 4b). These results suggest that NMI-induced neutrophil migration primarily occurs through the CXCR2 receptor.

**Fig. 4.**
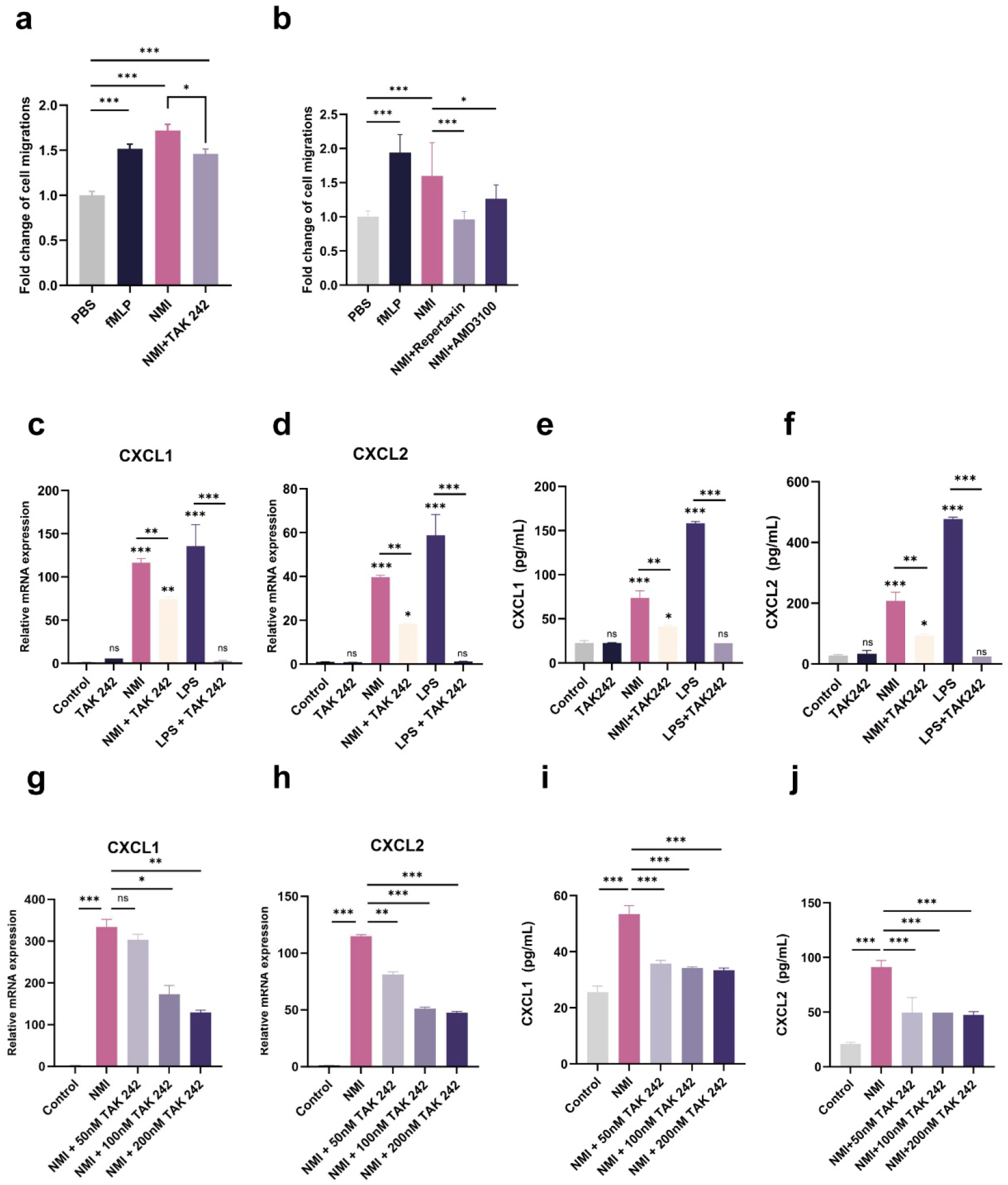
NMI causes cell migration by inducing neutrophils to release chemokines and is dependent on CXCR2. **a-b** In vitro transwell migration assay. The fold change of neutrophils migrations during through transwell filters toward fMLP (100 nM), and NMI (10 μg/mL). To inhibit the TLR4, CXCR2, CXCR4 receptor, neutrophils were pre-treated with TLR4 inhibitor TAK242(100 nM), CXCR2 inhibitor Repertaxin (1 μM) and CXCR4 inhibitor AMD3100(1 μM) for 30 min before NMI treatment. **c-J** Expression and secretion of chemokines after NMI stimulation of neutrophils. The **c, d, g** and **h** were qRT-PCR assay for transcriptome expression of CXCL1, CXCL2 after stimulation of neutrophils with 10 μg/mL NMI protein for 2 hours, and the **e, f, I** and **j** were ELISA assay for release of CXCL1, CXCL2 into supernatant levels, with the LPS group as a positive control. The inhibition group was incubated with TAK242 for 30 min before the start of the experiment, and the other groups were added with an equal amount of solvent. All data are expressed as mean ± SEM. Statistical significance was determined using one-way ANOVA (n = 3, *P< 0.05, **P< 0.01, ***P< 0.001).

CXCR2 serves as the receptor for several chemokines that play a crucial role in neutrophil migration [24]. We investigated the expression and release of CXCR2 ligands, specifically CXCL1 and CXCL2, from neutrophils that were directly stimulated in vitro with NMI proteins. As illustrated in Fig. 4c-f, treatment with 10 μg/mL NMI resulted in a significant increase in the expression and release of CXCL1 and CXCL2 into the supernatant. In alignment with the findings from the transwell assay, the LPS-treated group exhibited complete inhibition of chemokine expression and release at a concentration of 100 nM TAK242, while this inhibition was only partial in response to NMI. To rule out the possibility that the observed effects were solely due to the concentration of TAK242, we pre-treated the cells with varying concentrations of TAK242 (50 nM, 100 nM, and 200 nM) before stimulating them with the NMI recombinant protein. The results indicated that the inhibitory effect remained relatively unchanged with increasing TAK242 concentrations (Fig. 4g-j). These findings suggest that NMI regulates the expression of neutrophil chemokines CXCL1 and CXCL2 in part through its receptor TLR4, and this process may also engage additional regulatory pathways.

### 5. NMI acts on neutrophils to express CXCL1 through activation of the NF-κB signaling pathway

To assess the activation of signaling pathways following neutrophil stimulation with NMI, we conducted RNA-seq transcriptomics analysis on samples treated with 10 μg/mL NMI protein. As illustrated in Fig.5a, KEGG pathway enrichment analysis revealed that several inflammatory response-related pathways were significantly activated in comparison to the control group. Notably, genes associated with the NF-κB signaling pathway exhibited marked activation following stimulation with the NMI recombinant protein.

**Fig. 5.**
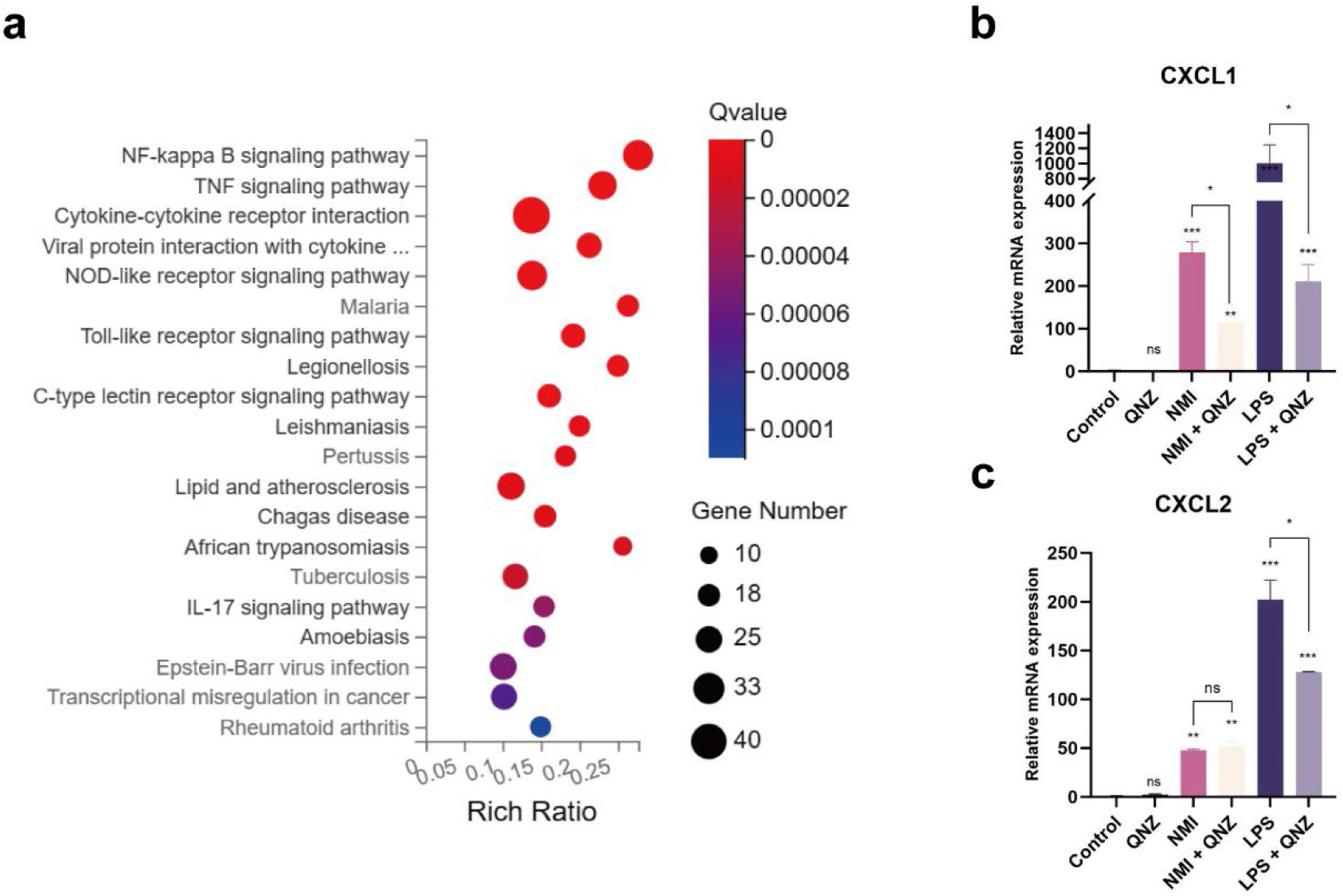
NMI acts on neutrophils to express chemokines through activation of the NF-κB pathway. **a** KEGG pathway enrichment analyses of the upregulated DEGS. Dot size represents the number of DEGS, and the dot color represents the corresponding Q value. **b** and **c** qPCR assay for transcriptome expression of CXCL1 and CXCL2 after stimulation of neutrophils with 10 μg/mL NMI recombinant protein for 2 hours, 1 μM QNZ added to the cells for 30 min of incubation before the experiment, and other groups were added with equal amount of solvent. All data are expressed as mean ± SEM. Statistical significance was determined using one-way ANOVA (n = 3, *P< 0.05, **P< 0.01, ***P< 0.001).

Since NMI is known to activate the NF-κB signaling pathway in mouse RAW264.7 cells [14], we investigated the activation of this pathway in neutrophils following stimulation with NMI protein. To assess the phosphorylation status of the NF-κB p65 protein, we employed immunoblotting techniques. After treating mouse neutrophils with 10 μg/mL of recombinant NMI for 15 min, we detected phosphorylation of the NF-κB p65 protein (Fig. S3). Furthermore, to validate that the NF-κB signaling pathway is responsible for the chemokine production following NMI stimulation, we pre-treated neutrophils with QNZ, a specific inhibitor of the NF-κB pathway, for 30 minutes prior to stimulation with 10 μg/mL of NMI for 2 hours. As illustrated in Fig. 4b-c, inhibition of the NF-κB signaling pathway resulted in a significant reduction in the expression of the chemokine CXCL1, while the expression of CXCL2 remained unaffected. These findings indicate that NMI triggers the NF-κB signaling pathway, leading to an increase in CXCL1 expression, which in turn facilitates neutrophil migration.

## Discussion

In this study, we elucidate the pivotal role of NMI in the recruitment of neutrophils, demonstrating its ability to attract a substantial number of these cells in vivo without being entirely reliant on macrophages. Furthermore, our findings indicate that NMI-stimulated neutrophils express and secrete a significant quantity of chemokines via the NF-κB signaling cascade. Notably, the inhibition of the receptor CXCR2 substantially mitigated NMI-induced neutrophil migration. In contrast, the inhibition of TLR4 and CXCR4 exhibited only a moderate inhibitory effect.

The recruitment of neutrophils is a complex process involving the interplay of multiple cell types, cytokines, and chemokines. Precise and directed migration to the site of inflammation is facilitated by chemokine guidance [25]. The IFP35 family protein NMI has been implicated as a DAMP involved in both acute and Chronic inflammatory diseases [14, 15, 26]. In our study, we aimed to elucidate the role of NMI in neutrophil recruitment by expressing and purifying the NMI protein. Our findings from both the air pouch model and the peritonitis model demonstrated that exogenous NMI effectively recruits neutrophils in vivo. Interestingly, knockout of *nmi* did not inhibit thioglycollate-induced neutrophil recruitment, suggesting that this particular pathway is independent of NMI. Moreover, we observed that the neutrophil recruitment induced by NMI was not entirely inhibited after the removal of macrophages. This finding implies that while NMI can activate macrophages to express large amounts of chemokines, which are the most potent chemical signals for neutrophil recruitment, there may also be other recruitment mechanisms at play. Previous studies have shown that other DAMPs, such as HMGB1 [27], S100A8/A9 [28] can chemically induce neutrophil recruitment during inflammatory responses. Consistent with this, our results from both transwell and agarose migration assays revealed that NMI also exerts a direct, chemically-induced effect on neutrophil recruitment in vitro.

Previous studies have demonstrated that DAMPs influence neutrophils through a variety of receptors and signaling pathways. For instance, Florence et al. demonstrated that elevated levels of HMGB1 can induce chemotaxis in human neutrophils by stimulating the production of IL-8. This process is mediated through the activation of TLR2, TLR4, and RAGE receptors [19]. Additionally, HMGB1 can also form a complex with CXCL12, which subsequently bind to CXCR4, promoting the recruitment of inflammatory cells to the site of tissue damage [29]. Research by Anna Moles et al. further indicated that TLR2 and S100A8/S100A9 are key regulatory factors for the expression of CXCL-2 and the recruitment of neutrophils in the liver [30]. In our study, we found that inhibiting TLR4 did not completely block the migration of neutrophils induced by NMI or the expression of chemokines, this finding contrasts with previous conclusions that TLR4 serves as a receptor for NMI in macrophages, suggesting that NMI may engage additional, yet unidentified, receptors and pathways in neutrophils. Furthermore, we observed that blocking CXCR2 effectively inhibited NMI-induced neutrophil migration. Although there is currently no direct evidence linking NMI to CXCR2, our results imply that the promotion of neutrophil migration by NMI is dependent on CXCR2. The identification of other potential receptors for NMI on neutrophils, as well as the specific interactions between NMI and CXCR2, warrants further investigation. In addition, the results of RNA-seq indicated that genes related to the NF-κB signaling pathway were heavily activated after NMI stimulation of neutrophils, highlighting this pathway as an important source for chemokines production. Finally, these activated and released chemokines induce the migration of neutrophils through the CXCR2 receptor.

However, the present study has several limitations. Firstly, while we employed a variety of pharmacological inhibitors to investigate neutrophil migration and chemokine secretion, we were unable to validate our findings with knockout cell models for critical genes such as TLR4 and CXCR2. Secondly, our understanding of NMI in the context of inflammation remains incomplete. Although we have confirmed that TLR4 serves as a receptor for NMI, our findings indicate the potential involvement of additional receptors. Therefore, further research is essential to clarify the role and mechanisms of NMI and its receptors in neutrophil recruitment.

In conclusion, our study uncovers a novel pathway through new DAMP—NMI on neutrophils recruitment. We demonstrated that NMI serves as a key regulator in the immune response, facilitating the recruitment of neutrophils to sites of inflammation and influencing the progression of inflammatory processes.

## Acknowledgements

We thank Prof. Huanhuan Liang (Sun Yat-sen University) for valuable discussion and critical comments on the manuscript. We also thank the School of Medicine of Sun Yatsen University and the Centre for Laboratory Animals of Sun Yat-sen University for providing the experimental site and instrumental support.

## List of abbreviations

NMI: N-myc and STAT interactor
IFP35: interferon-inducible protein 35
DAMP: Danger-associated molecular pattern
TLR4: toll-like receptor 4
CXCR2: CXC chemokine receptor 2
CXCR4: CXC chemokine receptor 4
PAMPs: pathogen-associated molecular patterns
PRRs: pattern recognizing receptors
TLRs: Toll-like receptors
RAGE: receptor for advanced glycation end products
HMGB1: high-mobility group box protein 1
S100A8/A9: Calprotectin

## Declarations

### Ethics approval and consent to participate

Not applicable.

### Consent for publication

Not applicable.

### Availability of data and materials

The data and materials used in this study are available from the corresponding authors upon reasonable request.

### Author contributions

Z.C. and Y.Y. designed and performed the experiments. Z.C. prepared the images. P.Y helped to performed the animal model building. Z.C., Z.W., N.X., and Y.L. analyzed the data, wrote the manuscript and organized the study. All authors contributed to the manuscript and approved the version as submitted.

