## Supplementary file 1 for "NMI induces chemokine release and recruits neutrophils through the activation of NF-κB pathway"

### Supplementary Figure

**Fig. S1**

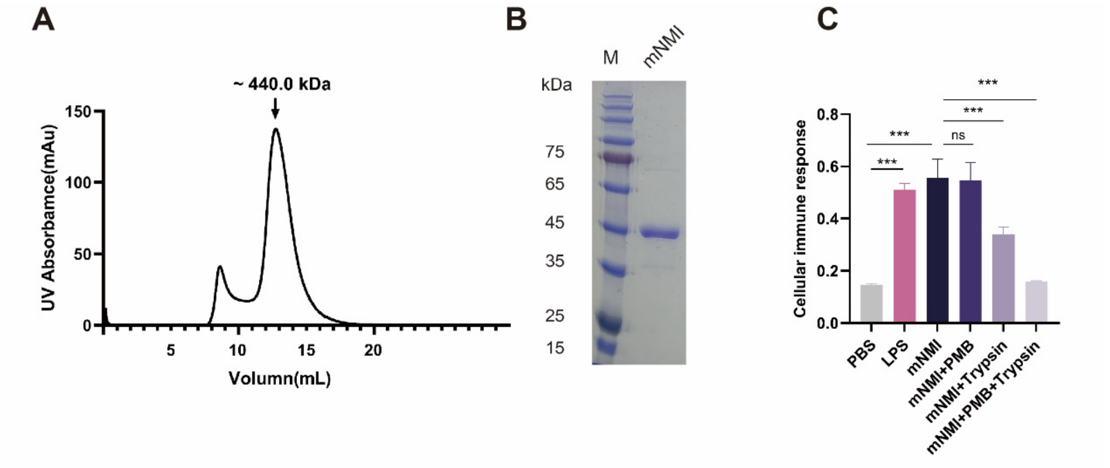


**Fig. S1 Purification, identification and protein activity assay of NMI recombinant proteins**

**(A)** Peak diagram of mNMI passing through Superdex200 10/300 GL column with PBS buffer.

**(B)** Verification of mNMI by SDS-PAGE and Caumas Brilliant Blue staining.

**(C)** Detection of cellular NF-κB pathway activation after stimulation of RAW-Blue[1] cells with PBS, mNMI (10 μg/mL), mNMI + PMB[2], mNMI + Trypsin, and mNMI + PMB + Trypsin.

**Fig. S2**

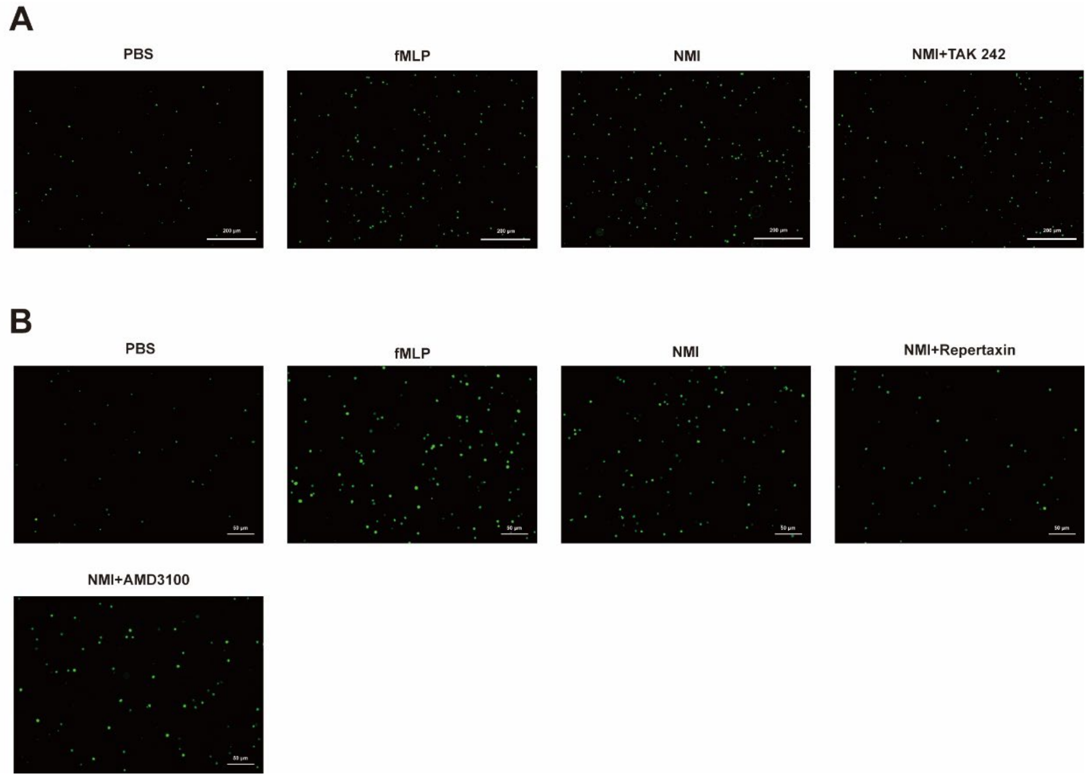


**Fig. S1 In vitro transwell migration assay.**

**A** Visualization of neutrophils chemotaxis with and without NMI stimulation (CFSE staining), neutrophils were pretreated with TLR4 inhibitor TAK242.

**B** Visualization of neutrophils chemotaxis with and without NMI stimulation (CFSE staining), neutrophils were pretreated with CXCR2 inhibitor Repertaxin or CXCR4 inhibitor AMD3100.

**Fig. S3**

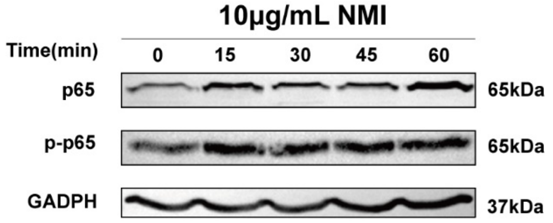


**Fig. S3** NMI stimulates neutrophils for different times (0 min, 15 min, 30 min, 45 min, 60 min) Phosphorylated and total p65 were evaluated with Western blot.
